# Characterization of *Vibrio cholerae* isolates from freshwater sources in northwest Ohio

**DOI:** 10.1101/2020.04.30.070565

**Authors:** Judy Daboul, Logan Weghorst, Cara DeAngelis, Sarah Plecha, Jessica Saul-McBeth, Jyl S. Matson

## Abstract

*Vibrio cholerae* is a natural inhabitant of aquatic ecosystems worldwide, typically residing in coastal or brackish water. While more than 200 serogroups have been identified, only serogroups O1 and O139 have been associated with epidemic cholera. However, infections other than cholera can be caused by nonepidemic, non-O1/non-O139 *V*. *cholerae* strains, including gastroenteritis and extraintestinal infections. While *V*. *cholerae* can also survive in freshwater, that is typically only observed in regions of the world where cholera is endemic. We recently isolated *V*. *cholerae* from several locations in lakes and rivers in northwest Ohio. These isolates were all found to be non-O1/non-O139 *V*. *cholerae* strains, that would not cause cholera. However, these isolates contained a variety of virulence genes, including *ctxA, rtxA, rtxC, hlyA*, and *ompU*. Therefore, it is possible that some of these isolates have the potential to cause gastroenteritis or other infections in humans. We also investigated the relative motility of the isolates and their ability to form biofilms as this is important for *V*. *cholerae* survival in the environment. We identified one isolate that forms very robust biofilms, up to 4x that of our laboratory control strains. Finally, we investigated the susceptibility of these isolates to a panel of antibiotics. We found that many of the isolates showed decreased susceptibility to some of the antibiotics tested, which could be of concern. While we do not know if these isolates are pathogenic to humans, increased surveillance to better understand the public health risk to the local community should be considered.

**Importance:** This study found that *Vibrio cholerae* belonging to non-O1/non-O139 serogroups is present in freshwater lakes and rivers in northwest Ohio. Although non-O1/non-O139 *V*. *cholerae* strains generally do not produce cholera toxin, and thus do not cause epidemic cholera, they often contain other virulence factors that may contribute to pathogenicity. In fact, we found that these local isolates contained an assortment of potential virulence genes. We also found that some of the isolates showed antibiotic resistance. While we do not currently know if the local *V*. *cholerae* strains are capable of causing disease in humans, increased monitoring for *V*. *cholerae* in the region is warranted in the interest of public health.

## Introduction

*Vibrio cholerae* is the bacterium responsible for the epidemic diarrheal disease cholera, but it is also a natural inhabitant of aquatic environments in many regions of the world (1). Due to their widespread existence in the environment, *Vibrio* species encounter a wide range of unfavorable conditions to which they have to sense and respond appropriately, by upregulating or downregulating specific gene expression. These conditions are also constantly changing due to natural and manmade events. Despite these challenges, due to its unique ability to adapt to and thrive in diverse and changing environments, *V*. *cholerae* has been a very successful marine organism and human pathogen. In turn, *V*. *cholerae* causes outbreaks of disease that are not always linked to predictable water sources. Therefore, it is critical that we better understand the varied locations where *V*. *cholerae* can survive worldwide in order to develop measures to limit human exposure to these ubiquitous aquatic bacteria.

*V*. *cholerae* strains are divided into more than 200 different serogroups based on O antigen structure. Historically, the O1 serogroup has been primarily responsible for causing epidemic cholera, and in 1992 another epidemic serogroup, O139, emerged (2). Pathogenic (toxigenic) *V*. *cholerae* strains encode two major virulence factors: cholera toxin (CT) and the toxin-coregulated pilus (TCP), the expression of which leads to the severe diarrhea associated with cholera disease (3). The remaining *V*. *cholerae* strains are generally referred to as “non-O1, non-O139” strains and do not cause cholera epidemics. These strains are also often referred to as “non-toxigenic” as they rarely carry the genes for CT and TCP. However, many of these non-O1/non-O139 strains encode other virulence genes, and have caused numerous diseases ranging from gastroenteritis to extraintestinal infections in humans (4-6). These accessory virulence factors include the repeats-in-toxin (RTX) toxin cluster (7, 8), outer membrane proteins (including OmpU) (9, 10), and hemolysins (11, 12). The presence of these virulence factors in non-O1/non-O139 strains are assumed to play some role in their survival in the natural environment (11). However, due to their likely role in the pathogenesis of non-O1/non-O139 strains, the presence of these accessory virulence factors is often monitored when characterizing *V*. *cholerae* strains isolated from environmental and food sources (11-14).

*V*. *cholerae* is typically found in estuarine or coastal environments, with most toxigenic strains residing in subtropical climates such as southeast Asia. *V*. *cholerae* is one of the only *Vibrio* species that can tolerate the absence of salt, and transmission of cholera from freshwater sources occurs, but typically in regions of the world where pathogenic strains are endemic (15). In the United States, *V*. *cholerae* can be isolated from coastal environments, such as the Chesapeake Bay (16). Typically, these North American strains are non-toxigenic, but studies have shown that there is a high degree of genetic diversity in the isolates and they contain a random assortment of virulence genes that may warrant increased surveillance (4).

In this study, we report that there are environmental *V*. *cholerae* strains present in freshwater lakes and rivers around the area of Toledo, Ohio. As most vibrios prefer at least a small amount of salt in their habitat and are typically found in warmer climates, this is unexpected. We have isolated *V*. *cholerae* from multiple locations over multiple dates in the Maumee river and along the shores of Lake Erie (Figure 1). These locations are all easily accessed by the local population and commonly used for fishing and recreational purposes. Many of these isolates were found in river water that should not be a part of shipping routes, so these strains are unlikely to have been recently introduced by expelled ballast water from large ships. Initial characterization showed that all of the isolates collected to date are from non-O1/non-O139 serogroups. The isolates were further examined for the presence of virulence genes, antibiotic resistance, motility, hemolysis activity, and biofilm formation. Based on the fact that we identified *V*. *cholerae* in water collected from most of these sampling locations, and that the isolates contain diverse phenotypic and genotypic traits, we hypothesize that *V*. *cholerae* is a normal member of the regional aquatic microbiota and it has been present in the area for some time.

**Figure 1.**
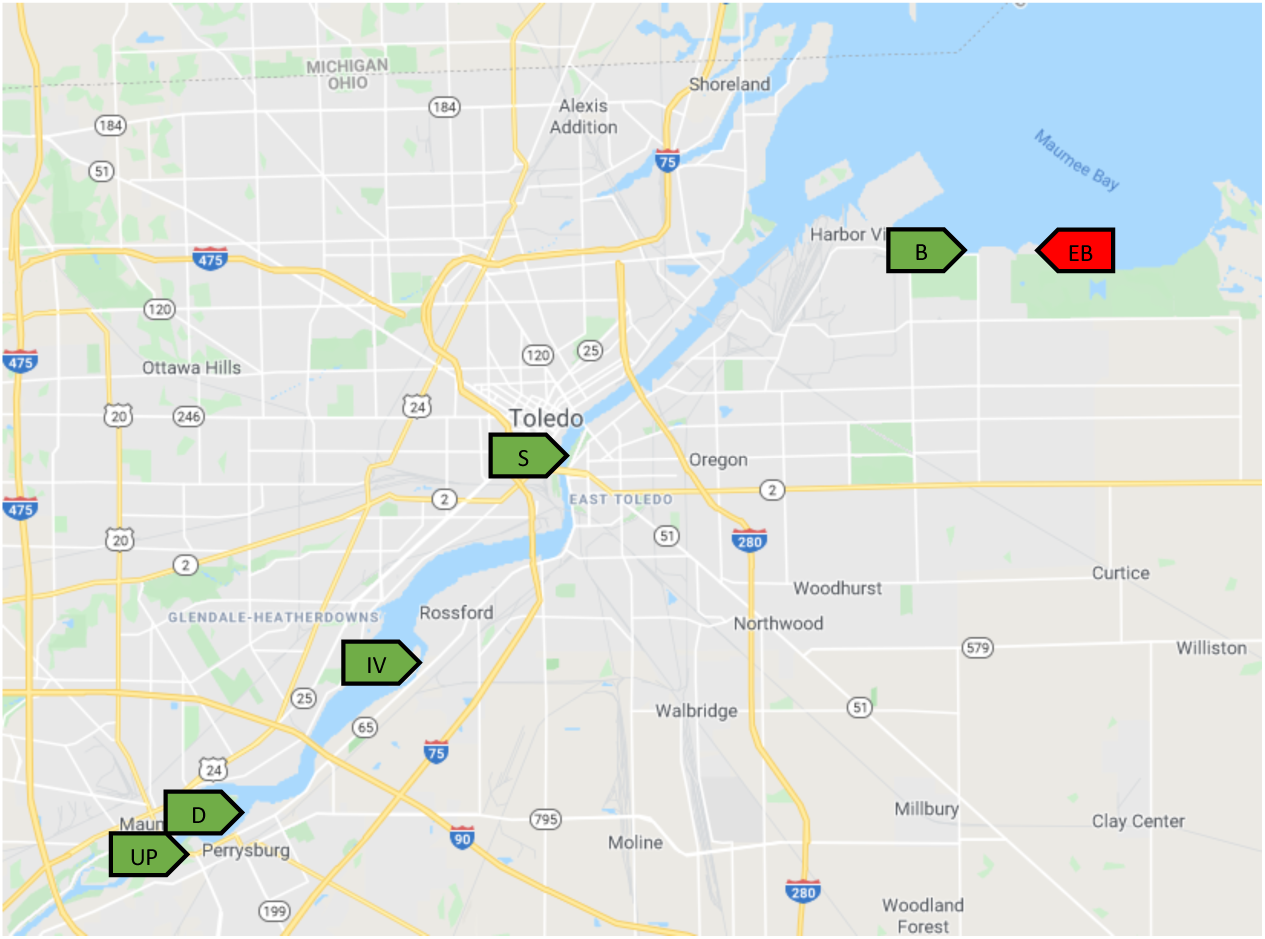
Water collection sites in the Toledo, Ohio area. The green markers indicate locations where *V*. *cholerae* was isolated from water samples. The red marker indicates a location that was negative for *V*. *cholerae*. Locations UP, D, and IV were sampled in both 2017 and 2018.

## Materials and Methods

### Sample collection and isolation of *Vibrio cholerae*

Water was collected during the summers of 2017 and 2018 from various locations on the Maumee River and the shore of Lake Erie (Figure 1). *V*. *cholerae* was isolated using alkaline peptone water enrichment according to standard protocols (17). Samples of water (∼250 mL) were collected and filtered using 0.22 μm pore size membrane filters. The membranes were then incubated statically in 100 mL alkaline peptone water (1% peptone, 1% NaCl, pH 8.6) overnight at 30°C. 100 μL of the culture (taken from the surface) was streaked onto thiosulfate citrate bile sucrose (TCBS) agar (BD Difco, USA) and incubated overnight at 37°C. Yellow colonies were isolated for species identification using PCR for the *V*. *cholerae* species-specific *ompW* gene (18). Antiserum kits for O1 and O139 *V*. *cholerae* (Hardy diagnostics, Santa Maria, CA) were used to determine serotype by slide agglutination, according to the manufacturer’s instructions.

### Growth and motility

Strains were routinely cultivated in LB medium at 37°C. Bacteria were also grown in M9 media to assess biofilm formation in minimal media. Bacterial isolates were maintained at −80°C in LB containing 20% glycerol. Hemolysis activity was investigated by growth of the isolates on Trypticase Soy Agar (TSA) with 5% sheep blood (BD). Motility assays were performed on LB media containing 0.3% agar, with total area of bacterial spread measured after 24 h of incubation at 37°C.

### PCR genotyping

Genomic DNA was extracted using the boiled template method (19). PCR was performed in a 25 μL reaction mixture using Radiant HiFi Ultra polymerase (Alkali Scientific Inc.). Gene targets and oligonucleotide sequences are listed in Table S1. *V*. *cholerae* reference strains C6706 and O395 were included as controls.

### Biofilm formation

Biofilm formation was measured using the crystal violet staining assay (20). Briefly, overnight cultures of the *V*. *cholerae* isolates were subcultured and grown statically in 24-well plates for the times indicated in either LB or M9 minimal media. After incubation, the culture media was removed and the wells were washed 3x with PBS. The wells were then stained with 1% crystal violet in 25% ethanol for 15 minutes and washed 3x with distilled water. The remaining crystal violet in the wells was solubilized using 33% acetic acid and the OD_570_ was measured. Biofilm quantity is represented as the OD_570_ divided by the OD_600_ of a duplicate culture.

### Antibiotic susceptibility

Antibiotic susceptibility was determined by disk diffusion on Mueller-Hinton agar (BD, USA) according to the Clinical and Laboratory Standards Institute guidelines for *V*. *cholerae* (21). The zone standards for *Enterobacteriaceae* were used when there were no established breakpoint interpretive criteria for *V*. *cholerae*. *V*. *cholerae* reference strains C6706 and O395 were included as controls. All strains were tested for susceptibility to tetracycline, 30 μg; trimethoprim-sulfamethoxazole, 1.25/23.75 μg; chloramphenicol, 30 μg; ciprofloxacin, 5 μg; erythromycin, 15 μg; nalidixic acid 30 μg; streptomycin, 10 μg; gentamicin 10 μg.

## Results

### Detection and isolation of *V*. *cholerae*

*V*. *cholerae* is a naturally occurring component of marine and estuarine costal microbiota in North America. However, there has been little investigation as to whether or not it is present in non-coastal areas in this part of the world. Most *Vibrio* species require salt for growth, with *V*. *cholerae* being an exception (22). Therefore, it is possible that *V*. *cholerae* could be present in freshwater lakes and rivers, although it is generally assumed to reside in more tropical climates. In order to determine if *V*. *cholerae* could be isolated from freshwater sources in northwest Ohio, we collected water samples during the summers of 2017 and 2018. Water was obtained from several locations in the Maumee River and along the shores of Lake Erie (Figure 1). The locations were accessible from the shoreline or boat docks in areas used for boating, fishing, or other recreational activities. *V*. *cholerae* was isolated using alkaline peptone water enrichment (17), and all isolates were confirmed using PCR for the *V*. *cholerae* species-specific *ompW* gene (18). We obtained *V*. *cholerae* isolates from 5 of the 6 sampling locations in 2017. Three of these locations were again sampled in 2018, and all three yielded *V*. *cholerae* in the second year as well. A total of 13 different *V*. *cholerae* isolates have been identified from these locations to date. In order to determine whether any of the isolates belonged to the epidemic O1 or O139 serogroups, we performed agglutination assays using O1- and O139-specific antisera. All of the isolates were found to be non-O1/non-O139 *V*. *cholerae* (data not shown).

### Distribution of virulence genes in regional *V. cholerae* isolates

Clinically significant infections other than cholera can be caused by non-O1/non-O139 *V*. *cholerae*, which are continually reported worldwide (4-6). Although non-O1/non-O139 *V*. *cholerae* strains typically do not produce the cholera toxin, they often contain other virulence genes that contribute to their pathogenicity. In order to determine the virulence potential of these isolates, we used PCR to identify the presence of known virulence genes. Genes that are usually associated with toxigenic *V*. *cholerae* strains include those encoding the cholera toxin (*ctxA, ctxB*) and genes of the CTX element (*ace, zot*). In addition, we also looked for the presence of genes encoding the toxin-coregulated pilus (*tcpA, tcpI*) in these isolates. As might be expected, we did not identify most of these virulence genes in our non-O1/non-O139 isolates (Table 1). However, two of our isolates were positive for *ctxA*, which was unexpected. As these strains were negative for *ctxB*, they would likely not produce a functional cholera toxin.

**Table 1:** Genotypes of environmental *V*. *cholerae* isolates

| Isolate | *ompW* | *ompU* | *ctxA* | *ctxB* | *rtxA* | *rtxC* | *hlyA*<br>Classical | *hlyA* El<br>Tor | *tcpA*<br>Classical | *tcpA* El<br>Tor/O139 | *tcpI* | *ace* | *st* | *zot* |
| --- | --- | --- | --- | --- | --- | --- | --- | --- | --- | --- | --- | --- | --- | --- |
| IV-1 | + | + | - | - | + | + | + | + | - | - | - | - | - | - |
| IV-3 | + | - | - | - | - | - | - | - | - | - | - | - | - | - |
| IV-5 | + | + | - | - | + | + | + | + | - | - | - | - | - | - |
| UP2 | + | + | - | - | + | + | + | + | - | - | - | - | - | - |
| UP4 | + | + | - | - | + | + | + | + | - | - | - | - | - | - |
| UP5 | + | - | + | - | - | - | - | - | - | - | - | - | - | - |
| S1 | + | - | - | - | + | + | - | - | - | - | - | - | - | - |
| S2 | + | - | - | - | + | + | - | - | - | - | - | - | - | - |
| D1 | + | + | - | - | + | + | + | + | - | - | - | - | - | - |
| D3 | + | + | - | - | + | + | + | + | - | - | - | - | - | - |
| D8 | + | - | + | - | + | + | - | - | - | - | - | - | - | - |
| B1 | + | + | - | - | + | + | + | + | - | - | - | - | - | - |
| B4 | + | - | - | - | + | + | + | + | - | - | - | - | - | - |
| C6706 | + | + | + | + | + | + | + | + | - | + | + | + | - | + |
| O395 | + | + | + | + | - | - | + | - | + | - | + | + | - | + |

These isolates were also tested for the presence of accessory virulence factors, including the RTX toxin (*rtxA, rtxC*), outer membrane protein, *ompU*, and hemolysin, *hlyA*. All but two of the isolates were positive for both of the RTX genes, indicating the potential that most of our isolates could produce a functional RTX toxin (Table 1). The gene encoding the outer membrane protein OmpU was found in 7 of the 13 isolates. The presence of both the Classical and El Tor versions of the *hlyA* hemolysin gene in these isolates was also tested (12). The sequence of the *hlyA* gene has been used to differentiate between the two biotypes of *V*. *cholerae* O1. The sequence of the Classical biotype has an 11 base pair deletion within the *hlyA* gene, compared to the El Tor biotype (23). Eight of the 13 strains were positive for both variants of the *hlyA* gene. Based on these genotyping results, we can conclude that we have isolated a number of unique *V*. *cholerae* strains so far, and that they contain different combinations of potential virulence factors.

### Phenotypic assays

In order to further characterize these environmental *V*. *cholerae* isolates, we performed other phenotypic assays. We examined the motility of the strains by inoculating each of them onto LB motility media and measuring the area of bacterial spread after 24 h of incubation. There was a dramatic range of bacterial motility observed for the isolates (Figure 2). Six of the isolates were significantly more motile than our laboratory strains. None of the strains showed reduced motility compared to O395 and C6706. The wide range of motility demonstrated by the individual isolates suggests that we have isolated a number of unique *V*. *cholerae* strains and not simply a clonal population that is present in a number of locations.

**Figure 2.**
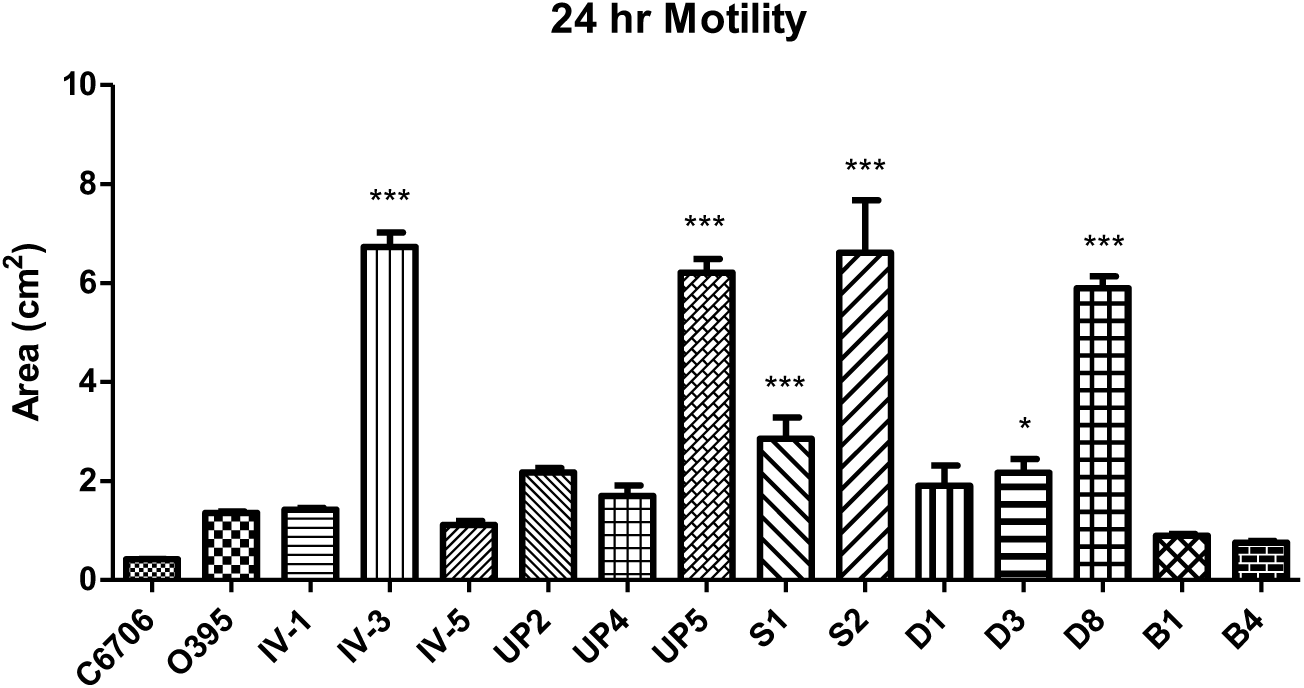
Motility of the *V*. *cholerae* isolates. A single colony of each isolate was inoculated into motility agar and incubated for 24 h. After incubation, the total area of bacterial spread was measured. The assay was performed in triplicate. Significance was compared to strain C6706, where *=p<0.5, **=p<0.1, ***=p<0.05.

We also examined the hemolytic patterns of each isolate by inoculation on 5% sheep blood nutrient agar plates and growing at 37°C overnight. All of the isolates demonstrated beta-hemolysis (data not shown). This was somewhat unexpected, as 5 of the isolates were negative by PCR for the presence of both the Classical and El Tor versions of the *hlyA* gene (Table 1). This suggests that many of our isolates encode and express another hemolysin other than the tested versions of *hlyA*.

### Isolate D1 produces elevated amounts of biofilm

When some of our isolates were cultured in liquid media (overnight, with shaking), we noticed differences in pellicle formation and/or sedimentation of the cultures (Figure 3A). Therefore, we hypothesized that there may be increased biofilm formation in some of the isolates as compared to our standard laboratory strains. To examine biofilm formation in these three isolates that routinely showed unusual growth characteristics, we performed crystal violet biofilm assays to determine the relative amounts of biofilm formed as compared to laboratory control strains. We first examined biofilm formation in LB media grown statically in polystyrene plates at two timepoints, after 18 and 24 h of growth. The El Tor strain C6706 forms significantly more biofilm compared to the Classical strain O395 at both timepoints (Figure 3B, C). When we compared the amount of biofilm formation in these three isolates to strain C6706, isolates IV-1 and D1 formed significantly increased amounts of biofilm at both timepoints, while isolate UP-2 formed less. Isolate D1 showed the most dramatic biofilm formation, producing approximately 4 times the amount of C6706 after 24 h of growth (Figure 3C). We also examined biofilm formation of these isolates when grown in minimal M9 media in order to determine if the elevated biofilm formation was dependent on the abundance of nutrients found in rich media. All strains show decreased growth in minimal media, resulting in the overall smaller ratios in OD_570_/OD_600_ (Figure 3D, E). However, despite this overall decreased growth, again isolate D1 formed significantly increased biofilms compared to strain C6706 at both timepoints. Due to the unusually high biofilm formation demonstrated by this strain, we had the genome sequenced (24). Sequence information can be found at DDBJ/ENA/GenBank (accession number QZCU00000000).

**Figure 3.**
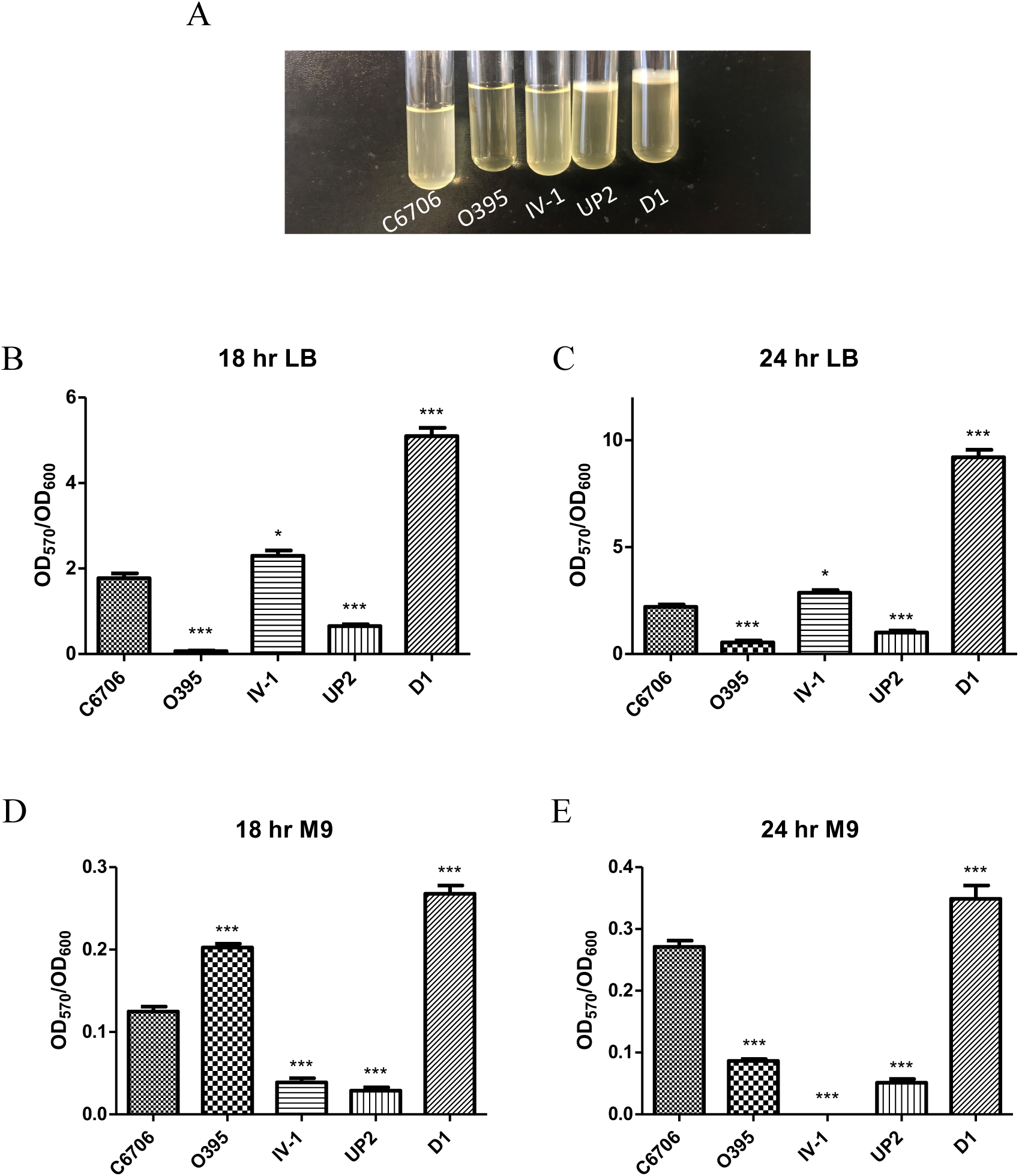
Biofilm formation of the environmental *V*. *cholerae* isolates. (A) Pellicle formation of selected environmental isolates compared to laboratory *V*. *cholerae* strains. The crystal violet assay was performed to determine biofilm formation after 18 h (B) or 24 h (C) of static growth in LB medium, and after 18 h (D) or 24 h (E) of static growth in M9 minimal medium. The assay was performed in triplicate. Significance was compared to strain C6706, where *=p<0.5, **=p<0.1, ***=p<0.05.

### Antibiotic resistance profiles of the environmental isolates

A major concern for any bacterial strains that may have public health consequences is the potential of resistance to commonly used antibiotics. Therefore, we performed antimicrobial susceptibility testing on our environmental isolates using a disk diffusion assay for 8 antibiotics. All strains were uniformly susceptible to chloramphenicol, tetracycline, trimethoprim-sulfamethoxazole, ciprofloxacin, and gentamicin (Table 2). Three of the isolates were found to be resistant to erythromycin. In addition, three isolates showed intermediate resistance to nalidixic acid and 6 others showed intermediate resistance to streptomycin (the laboratory strains have been selected for streptomycin resistance). Overall, while the isolates were sensitive to the majority of the antibiotics tested, it is worth noting that only 3 of the 13 isolates were fully susceptible to this relatively small panel of antibiotics (Table 2). Therefore, increased antimicrobial susceptibility testing may be warranted for these, and any future isolates collected from the area.

**Table 2:**
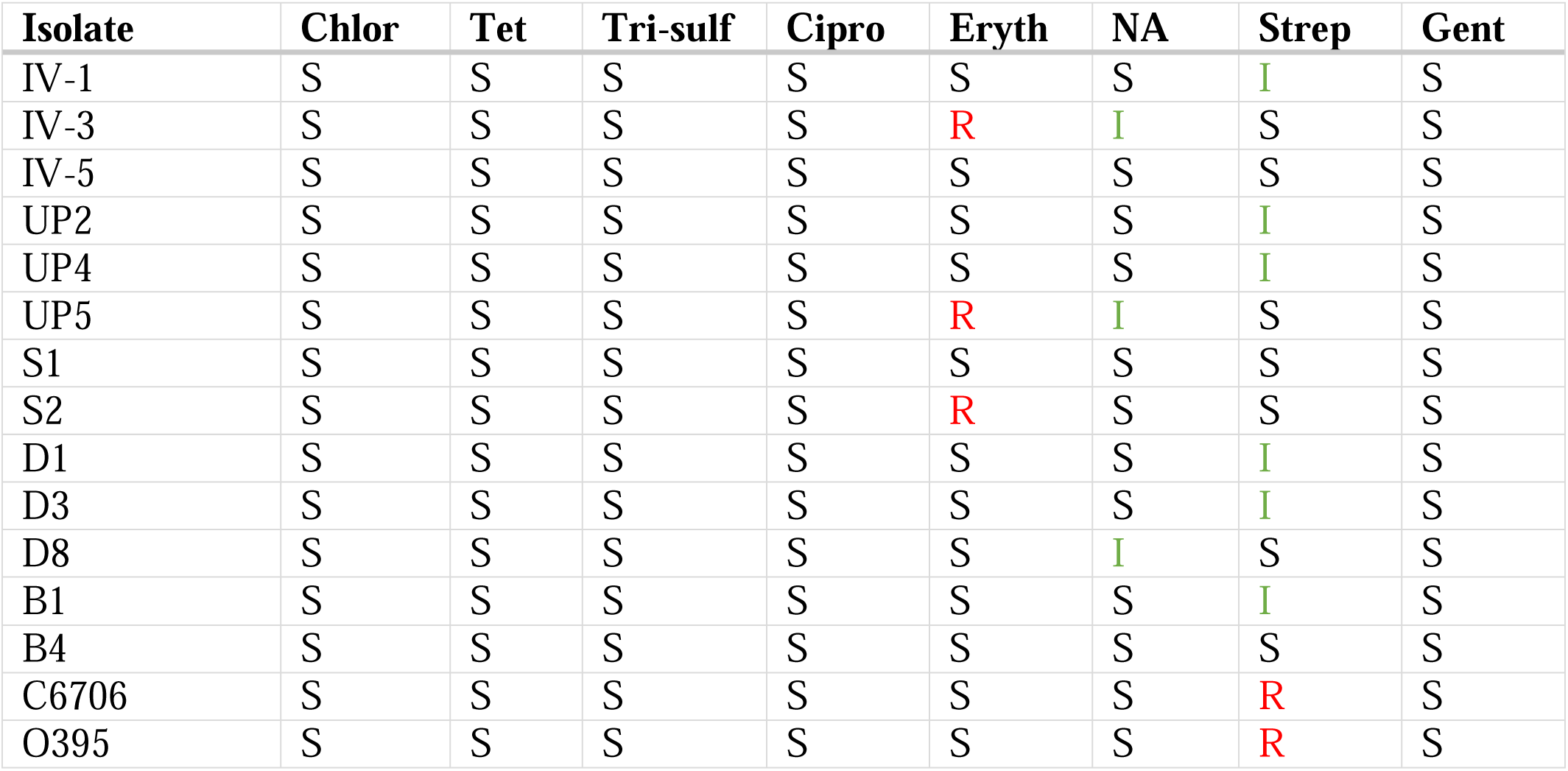
Antibiotic resistance profiles of *V*. *cholerae* isolates

| Isolate | Chlor | Tet | Tri-sulf | Cipro | Eryth | NA | Strep | Gent |
| --- | --- | --- | --- | --- | --- | --- | --- | --- |
| IV-1 | S | S | S | S | S | S | I | S |
| IV-3 | S | S | S | S | R | I | S | S |
| IV-5 | S | S | S | S | S | S | S | S |
| UP2 | S | S | S | S | S | S | I | S |
| UP4 | S | S | S | S | S | S | I | S |
| UP5 | S | S | S | S | R | I | S | S |
| S1 | S | S | S | S | S | S | S | S |
| S2 | S | S | S | S | R | S | S | S |
| D1 | S | S | S | S | S | S | I | S |
| D3 | S | S | S | S | S | S | I | S |
| D8 | S | S | S | S | S | I | S | S |
| B1 | S | S | S | S | S | S | I | S |
| B4 | S | S | S | S | S | S | S | S |
| C6706 | S | S | S | S | S | S | R | S |
| O395 | S | S | S | S | S | S | R | S |

## Discussion

The goal of this present study was to understand the presence, virulence, and antibiotic resistance profiles of *V*. *cholerae* in northwest Ohio. Although our sampling to date is relatively limited, it is clear that these bacteria are widely distributed in the local aquatic environment, being found in water from both lakes and rivers. While we have not estimated total abundance of vibrios in any given location at this time, *V*. *cholerae* has been readily isolated from relatively small samples of water in most of the locations sampled, suggesting it may be present in significant quantities. We have only identified non-O1/non-O139 *V*. *cholerae* to date, however these strains are more readily isolated in culture (25). *V*. *cholerae* O1 strains are known to be difficult to isolate, even in areas where cholera is endemic. Future studies will investigate the possibility that O1/O139 strains are also present in northwest Ohio using non-culture-based methods, such as the direct fluorescent antibody assay (26). This technique has been used to successfully detect O1 isolates in the Chesapeake Bay and other locations (13).

Previous studies have reported the presence of *V*. *cholerae* in freshwater samples in areas of the world that are not considered to be endemic for cholera. For example, in some lakes in Germany and Austria, pathogenic non-O1/non-O139 *V*. *cholerae* are present and occasionally cause human infections (5, 27). Only recently have researchers started exploring the possibility that *V*. *cholerae* may be present in non-coastal regions of North America. In addition to our studies, *V*. *cholerae* has been recently isolated from a freshwater lake in southwestern Pennsylvania (28). No studies to date have determined if *V*. *cholerae* isolated from these freshwater sources have demonstrated increased resistance to low temperatures or low salinities compared to isolates found in other regions of the world. Even if some of these strains show adaptation to cold weather, global climate change is expected to increase the surface water temperatures of lakes and rivers worldwide. This may lead to increased levels of vibrios in any water sources they are introduced, and thus, increased human infections. Therefore, it may be of public health benefit to institute increased monitoring programs for *V*. *cholerae* in these regions and inform clinicians to the possibility of screening for vibriosis, which may be overlooked as a possible cause of disease in local patients.

This study revealed that the freshwater non-O1/non-O139 *V*. *cholerae* isolated in northwest Ohio contain potential virulence genes. While *ctxA* was present in two of our isolates, the *ctxB* gene was not found in these strains. Based on the absence of *ctxB* and the fact that environmental non-O1/non-O139 isolates generally do not produce cholera toxin, we assume that these isolates would also not be able to produce a functional cholera toxin. However, vibriophages have been demonstrated to transfer toxin genes from CTXΦ-positive strains to environmental non-O1/non-O139 *V*. *cholerae* isolates (29). Again, future studies will involve more rigorous screening for O1 strains that could serve as a source of the *ctxA* gene found in some of our isolates.

Similar to numerous other studies examining non-O1/non-O139 *V*. *cholerae* in environmental, clinical, and seafood samples, we found that these isolates frequently contained genes encoding accessory virulence factors, including the RTX toxin, OmpU, and hemolysin. In *V*. *cholerae*, the *rtx* gene cluster encodes a multifunctional-autoprocessing RTX toxin (MARTX). This subgroup of RTX toxins are typically very large proteins (3500 to 5300 amino acids in length) that have multiple activities. The primary role of this toxin is evasion of the innate immune defenses during intestinal infection. The RTX toxin works in concert with HlyA, a hemolysin that was also detected in the majority of our isolates. Purified HlyA induces fluid accumulation and histological changes in the mucosa when injected into rabbit ileal loops (30). HlyA has also been shown to induce cell vacuolation and apoptosis in cultured mammalian cells (31, 32). *V*. *cholerae* strains deleted for both *rtxA* and *hlyA* are avirulent and unable to colonize mice (33). These toxins act together to inhibit clearance of the infecting bacteria from the small intestine by neutrophils, which is thought to explain their requirement for bacterial colonization (34). OmpU, one of the most abundant outer membrane proteins of *V*. *cholerae*, is required for bacterial resistance to antimicrobial peptides and induces IL-8 expression in intestinal epithelial cells, thus playing an important role in innate immunity (10, 35). The combined action of these virulence factors in these isolates of *V*. *cholerae* may enable these bacteria to induce gastroenteritis, or other infections, if ingested by a human.

In summary, we have found *V*. *cholerae* in freshwater lakes and rivers around Toledo, Ohio. While we do not have any evidence that these bacteria can cause human disease, we did identify several potential virulence genes in most of the isolates. In addition, several of the isolates showed some degree of antibiotic resistance. Therefore, increased monitoring for *V*. *cholerae* in the region, as well as increased characterization of the isolates, is warranted. Future studies aim to further characterize the relative resistance of these isolates to cold temperature and low salinity. We also would like to determine if these bacteria can be found in the intestines of local fish populations, as *V*. *cholerae* is known to colonize several fish species. This would be important information for regional commercial and recreational fishers. In addition, we also aim to sequence additional *V*. *cholerae* isolates from the area in order to identify other unexpected virulence factors and to determine the relationship of these isolates to sequenced environmental strains from other parts of the world.

## Acknowledgements

This study was supported by startup funds from the University of Toledo. JD and LW were supported by funds from the UT Medical Student Research Program. The authors would like to thank Laura Stanbery, Michael Nejedlik, and Travis Taylor for assistance in sample collection.

