## Supplementary material for "Characterization of *Vibrio cholerae* isolates from freshwater sources in northwest Ohio": Table S1

| **Primer** | **Sequence (5’-3’)** |
| --- | --- |
| *ompW* F | CACCAAGAAGGTGACTTTATTGTG |
| *ompW* R | GAACTTATAACCACCCGCG |
| *ompU* F | ACGCTGACGGAATCAACCAAAG |
| *ompU* R | GCGGAAGTTTGGCTTGAAGTAG |
| *ctxA* F | CGGGCAGATTCTAGACCTCCTG |
| *ctxA* R | CGATGATCTTGGAGCATTCCCAC |
| *ctxB* F | ATGCACATGGAACACCTCAAAATATTACTG |
| *ctxB* R | TCCTCAGGGTATCCTTCATCCTTTCAATC |
| *rtxA* F | CTGAATATGAGTGGGTGACTTACG |
| *rtxA* R | GTGTATTGTTCGATATCCGCTACG |
| *rxtC* F | TGCAAATCTCACATTAGCGCA |
| *rtxC* R | CCACTGCACCTTTCGGATACA |
| *hlyA* F (classical) | GGCAAACAGCGAAACAAATACC |
| *hlyA* F (El Tor) | GAGCCGGCATTCATCTGAAT |
| *hlyA* R | CTCAGCGGGCTAATACGGTTTA |
| *tcpA* F | CACGATAAGAAAACCGGTCAAGAG |
| *tcpA* R (classical) | TTACCAAATGCAACGCCGAATG |
| *tcpA* R (El Tor/O139) | CGAAAGCACCTTCTTTCACACGTTG |
| *tcpI* F | TAGCCTTAGTTCTCAGCAGGCA |
| *tcpI* R | GGCAATAGTGTCGAGCTCGTTA |
| *ace* F | TAAGGATGTGCTTATGATGGACACCC |
| *ace* R | CGTGATGAATAAAGATACTCATAGG |
| *st* F | GAGAAACCTATTCATTGC |
| *st* R | GCAAGCTGGATTGCAAC |
| *zot* F | TCGCTTAACGATGGCGCGTTTT |
| *zot* R | AACCCCGTTTCACTTCTACCCA |
